## Supplemental Information for "Non-Invasive Skin Sampling Detects Systemically Administered Drugs in Humans"

### Supplemental Methods

##### MZmine Parameters

MZmine version 2.37 was used to complete feature detection on the untargeted metabolomics data. The parameters are as follows: Mass Detection (MS1 noise level 5.0e4, MS2 noise level 1.0e3); ADAP Chromatogram Builder (MS level 1, min group size # of scans 5, group intensity threshold 5.0e4, min highest intensity 1.5e5, and m/z tolerance 0.001 m/z or 10 ppm); Chromatogram Deconvolution (Local Min Search, chromatographic threshold 1.00%, search minimum in RT range 0.2 min, minimum relative height 1.00%, minimum absolute height 1.5e5, min ratio of peak top/edge 1.5, peak duration 0.01 - 0.50 min, m/z center calculation median, m/z range for MS2 scan pairing 0.01 Da, RT range for MS2 scan 0.15 min); Isotope Peak Grouper (m/z tolerance 0.001 m/z or 5 ppm, retention time tolerance 0.15 min, maximum charge 3, representative isotope most intense); Join Aligner (m/z tolerance 0.001 m/z or 10 ppm, weight for m/z 75, retention time tolerance 0.15 min, weight for RT 25); Gap Filling (intensity tolerance 20%, m/z tolerance 0.001 m/z or 5 ppm, retention time tolerance 0.15 min); Peak Filter (area 2.0e4 - 1.0e20). The resulting feature tables were exported as a quantification table (.csv) and a spectral information table (.mgf).

##### Annotation Discussion

Note, diphenhydramine and diphenhydramine metabolites discussed were annotated based on exact mass, isotope pattern, and MS/MS spectral interpretation .

The molecular formula of diphenhydramine is $C_{17}H_{21}NO$ with a corresponding monoisotopic mass of 255.1623. The precursor ion observed for diphenhydramine was the proton adduct (i.e. $[M+H]^+$) with a monoisotopic *m/z* of 256.169. The mass error between the theoretical monoisotopic *m/z* and the measured monoisotopic *m/z* was -2.342 ppm. The observed isotope pattern (256.1698, 257.1733, 258.1769) matched that of the theoretical isotope pattern (https://www.lfd.uci.edu/~gohlke/molmass/?q=C17H22NO ). The MS2 consensus spectra from GNPS that was annotated as diphenhydramine had a precursor *m/z* of 256.1694 and mean retention time of 3.41 min (cluster 536). An authentic standard was analyzed and was observed at a retention time of 3.41 min.

*N*-desmethyldiphenhydramine has a molecular formula of $C_{16}H_{19}NO$ and a monoisotopic mass of 241.1467. The precursor ion observed for *N*-Desmethyldiphenhydramine was the proton adduct with a monoisotopic *m/z* of 242.154 which matches the theoretical monoisotopic *m/z* giving a mass error of 0 ppm. The annotated *N*-desmethyldiphenhydramine metabolite's MS2 consensus spectra from GNPS had a precursor *m/z* of 242.1540 and mean retention time 3.33 min (cluster index 484). The molecular formula of diphenhydramine *N*-oxide is $C_{17}H_{21}NO_2$ with a monoisotopic mass of 271.1572. The precursor ion observed for this metabolite was the proton adduct with a monoisotopic *m/z* of 272.165. The mass error between the theoretical monoisotopic *m/z* and the measured monoisotopic *m/z* was 1.837 ppm. The MS2 consensus spectra from GNPS annotated as diphenhydramine *N*-oxide had a precursor m/z of 272.1648 and mean retention time of 3.55 min (cluster 432). Diphenhydramine *N*-glucuronide has a molecular formula of $C_{23}H_{29}NO_7$ and a monoisotopic mass of 431.1944. The precursor ion observed for diphenhydramine was the proton adduct with a monoisotopic *m/z* of 432.201 with a mass error of -1.620 from the theoretical monoisotopic *m/z.* The MS2 consensus spectra from GNPS that was annotated as diphenhydramine *N*-glucuronide had a precursor *m/z* of 432.2013 and a mean retention time of 3.18 min (cluster 415).

The molecular formula of diphenhydramine *N*-glucose is $C_{23}H_{32}NO_6^+$ and has a monoisotopic mass of 418.223. The precursor ion observed for diphenhydramine *N*-glucose was the $[M]^+$ ion with a monoisotopic *m/z* of 418.223. The mass error between the theoretical monoisotopic *m/z* and the measured monoisotopic *m/z* was 1.195 ppm. The observed isotope pattern (418.2227, 419.2260, 420.2286) matches with the theoretical isotope pattern (https://www.lfd.uci.edu/~gohlke/molmass/?q=C23H32NO6%2B+). The MS2 consensus spectra from GNPS that was annotated as diphenhydramine *N*-glucose had a precursor *m/z* of 256.1694 and mean retention time of 3.18 min (cluster 794). An authentic standard was synthesized and analyzed and was observed at a retention time of 3.18 min.

**Three-Dimensional Spatial Model**

Spatial mapping was used to visualize diphenhydramine and diphenhydramine related metabolites detected on the skin. The normalized peak area median values were mapped onto the corresponding skin sites for each metabolite and displayed over the corresponding time course from the study.

Spatial mapping on the head and body model was generated using the ili surface mapping tool.(`ili: Spatial Data Mapping ) The 3D partial body model (obj file) was obtained from a free online 3D model platform.(Free 3D Models, Download or Edit Onli...) The 3D head model (obj file) was obtained from Sketchfab (Head model of a child by usd95). To map metabolite features onto a 3D model obj file, position coordinates (x,y,z and radius) must be assigned prior to mapping features with the ili tool. The assignment of skin site position coordinates for each 3D model obj was performed in Meshlab.(MeshLab ) These skin sites were approximate and representative of where the skin swab samples were collected for this study. A .csv file containing peak area median values and skin site specific coordinates and radii were dragged and dropped into ili along with the 3D model obj file to map and visualize the diphenhydramine and diphenhydramine related metabolites. The "White-Blue" color map was used to visualize the intensity of diphenhydramine and diphenhydramine related metabolites detected on each of the different skin sites over time.

| Subject_ID | AUCINF_obs (h*ng/mL) | Cl_F_obs (L/h) | Cmax (ng/mL) | Tmax (h) | HL (h) | Vz_F_obs (L) |
|---|---|---|---|---|---|---|
| 191001 | 2453.02 | 20.38 | 253 | 4 | 7.71 | 226.67 |
| 191002 | 2003.28 | 24.96 | 252 | 1.5 | 8.04 | 289.48 |
| 191003 | 1219.48 | 41 | 146 | 2 | 8.23 | 486.78 |
| 191004 | 849.76 | 58.84 | 85.9 | 2 | 8.19 | 695.49 |
| 191005 | 1222.65 | 40.89 | 146 | 2 | 8.24 | 486.14 |
| 191007 | 1647.45 | 30.35 | 252 | 2 | 7.38 | 323.24 |
| 191008 | 1498.64 | 33.36 | 138 | 1.5 | 11 | 529.65 |
| 191009 | 1234.67 | 40.5 | 141 | 2 | 8.13 | 475.22 |
| 191010 | 610.15 | 81.95 | 59.9 | 2 | 9.17 | 1084.4 |
| 191011 | 1007.84 | 49.61 | 102 | 2 | 8.77 | 627.7 |
| **Mean** | **1374.694** | **42.185** | **157.58** | **2.1** | **8.487** | **522.478** |
| Median | 1228.66 | 40.7 | 143.5 | 2 | 8.21 | 486.46 |
| SD | 548.651 | 17.98 | 71.204 | 0.699 | 1.014 | 245.658 |
| SE | 173.499 | 5.686 | 22.517 | 0.221 | 0.321 | 77.684 |

**Supplemental Table 1** Pharmacokinetic parameters for the 10 subjects were calculated as described above.

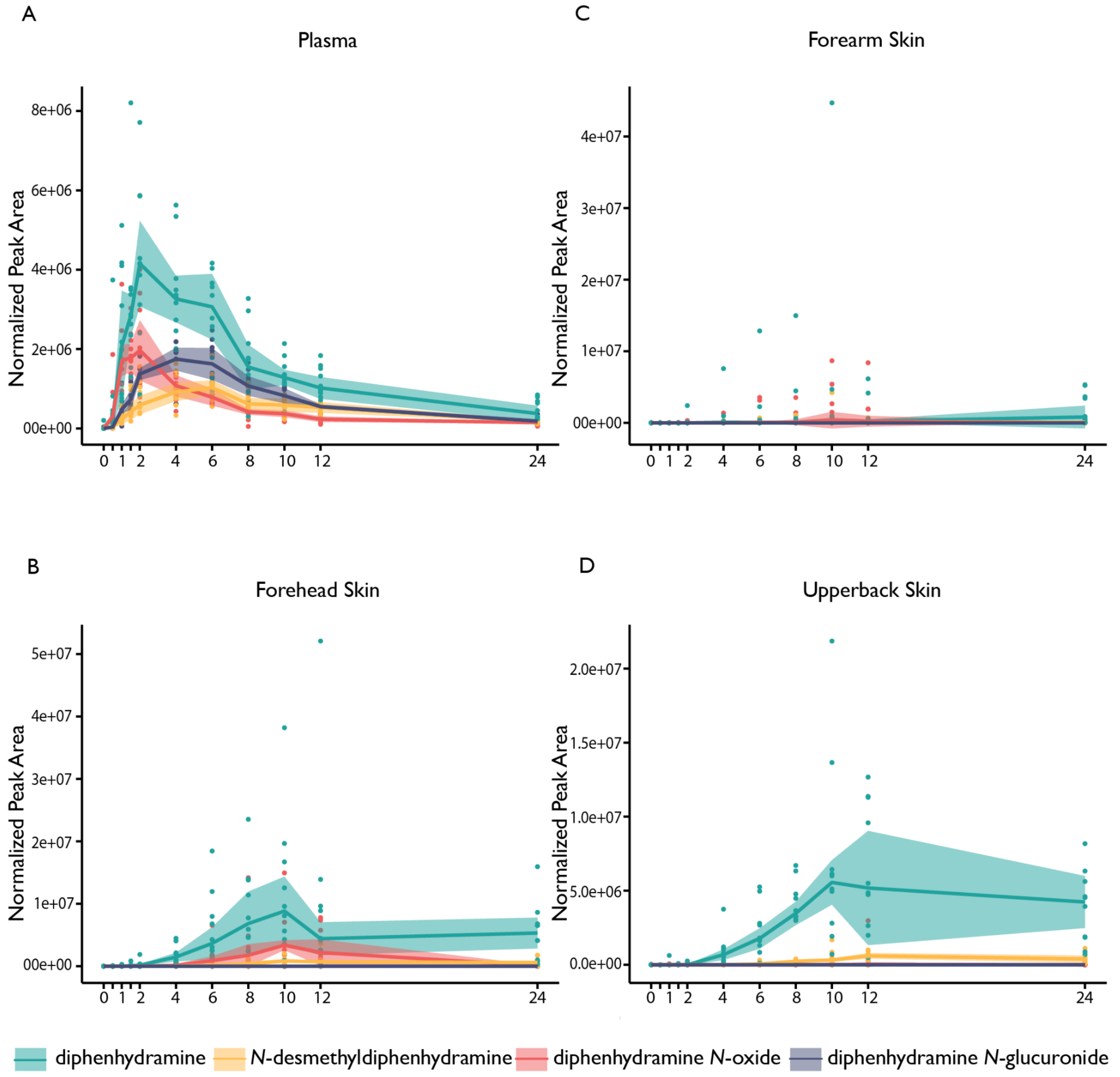


**Figure S1. Time versus peak area of diphenhydramine and diphenhydramine metabolites observed by sample type** Plot of time vs. peak area for diphenhydramine, *N*-desmethyldiphenhydramine, diphenhydramine *N*-oxide and diphenhydramine *N*-glucuronide in (**A**) plasma, (**B**) forehead skin, (**C**) forearm skin, and (**D**) upper back skin. The highlighted portion of these plots represent the interquartile range for each sample type and the solid line represents the median values.

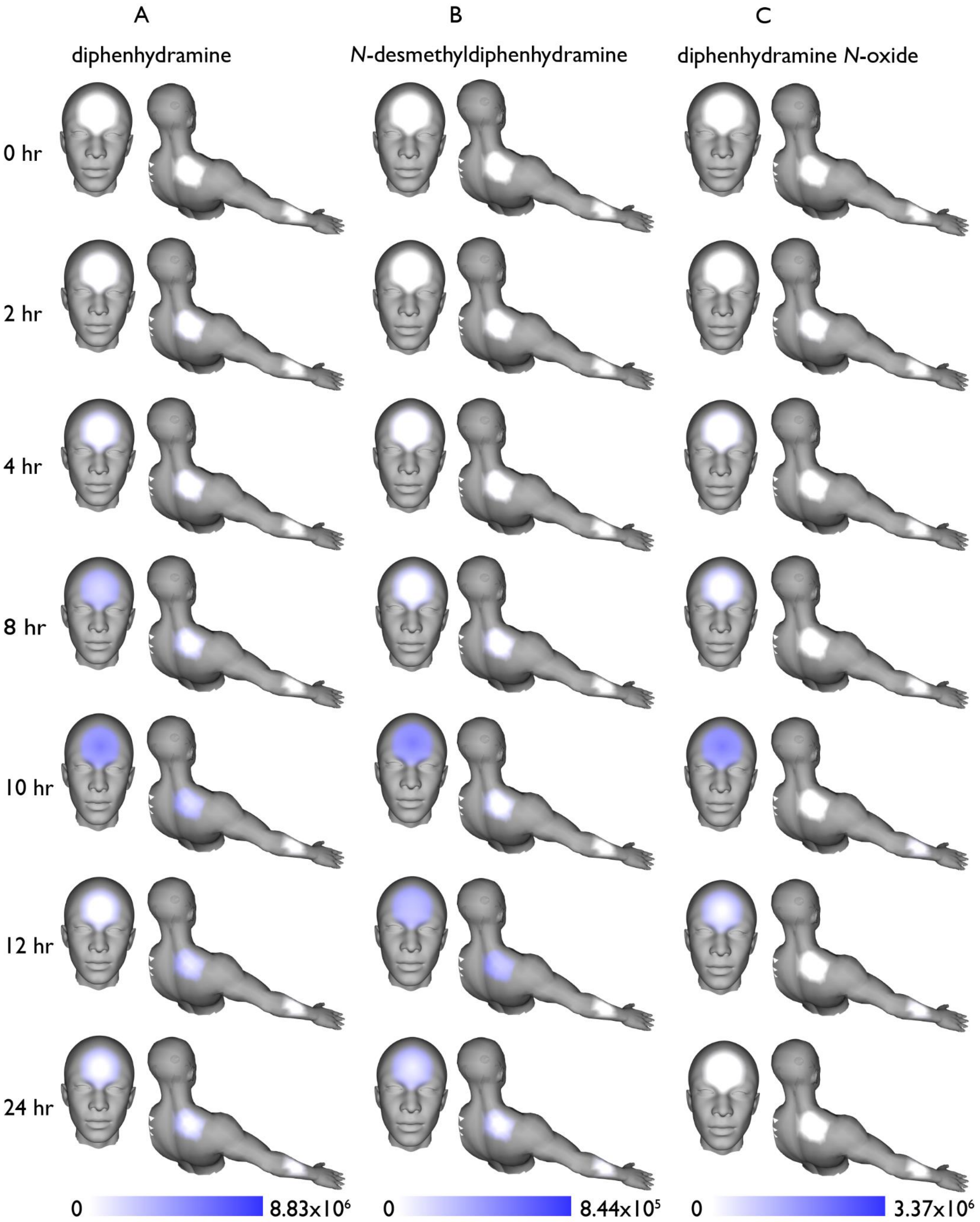


**Figure S2. Visualization of diphenydarmine and annotated metabolites on an androgynous model displaying median values over time.** This 3D illustrative molecular map uses a white-blue color scale representative of increasing metabolite intensity for each skin site observed for each metabolite (**A**) diphenhydramine, (**B**) *N*-desmethyldiphenhydramine and (**C**) diphenhydramine *N*-oxide.

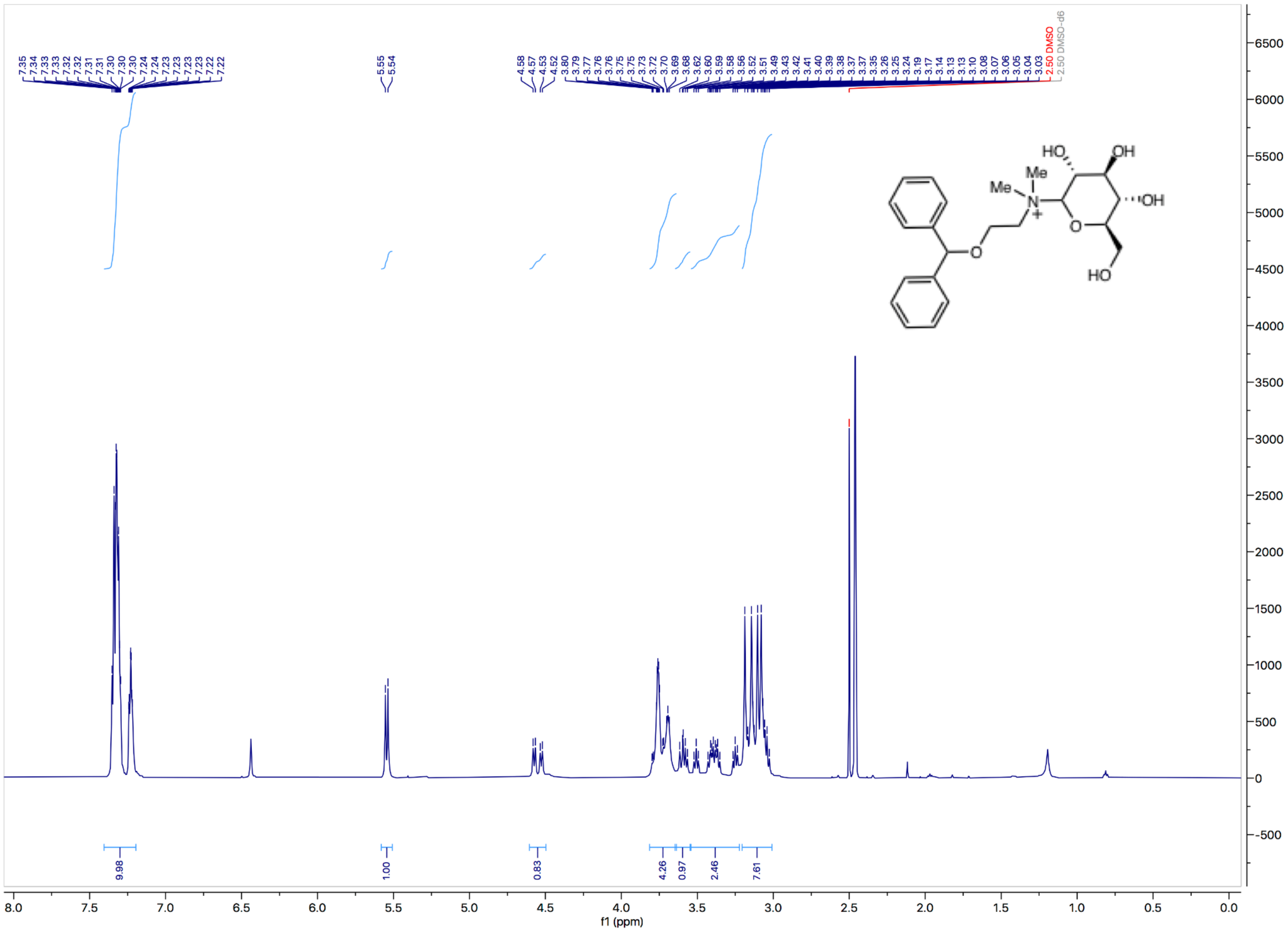

**Figure S3.**: $^1$H NMR of synthesized diphenhydramine-glucose metabolite.

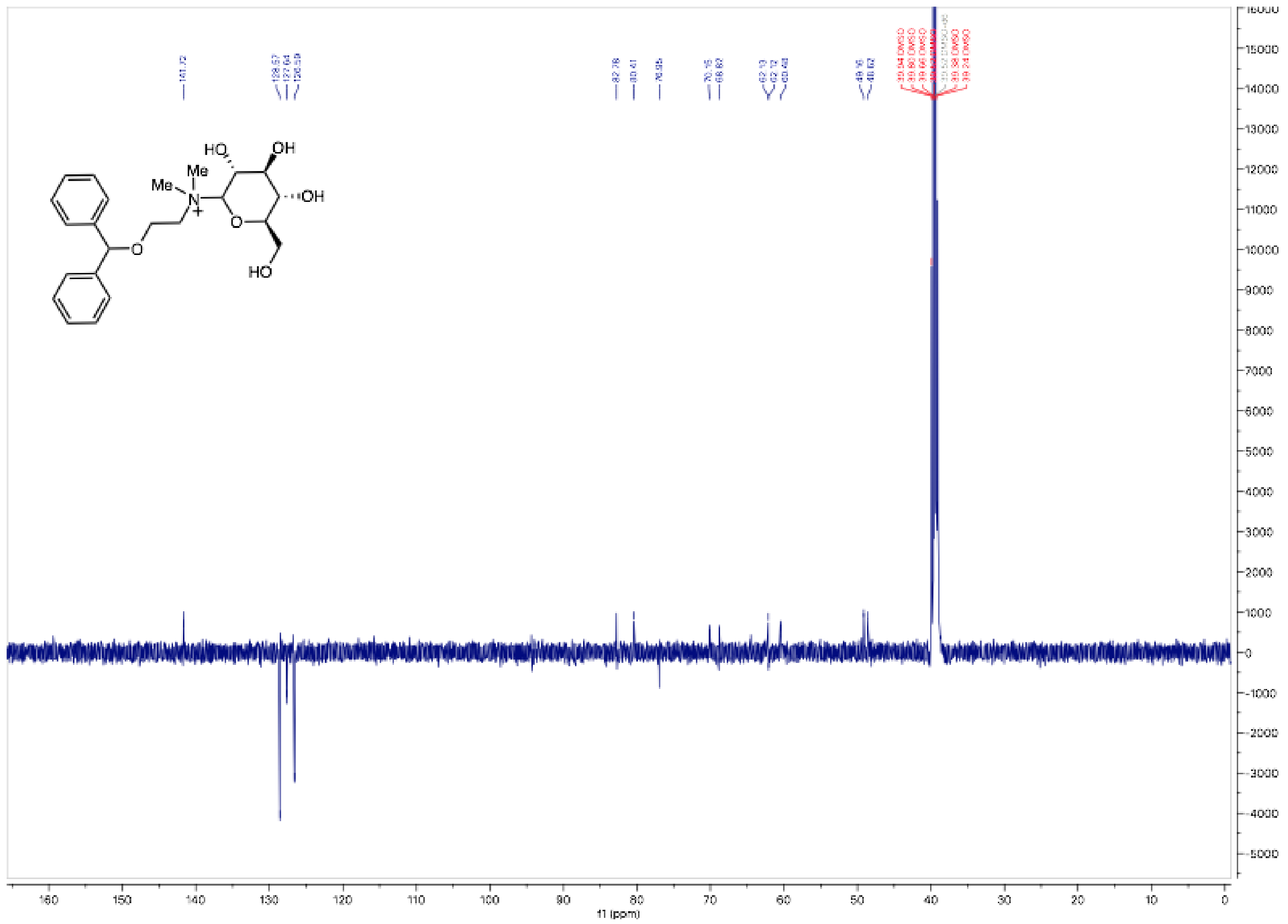

**Fig. S4**: $^{13}$C NMR of synthesized diphenhydramine-glucose metabolite in DMSO-$d_6$.
